# Chikungunya-virus-specific CD4^+^ T cells are associated with chronic chikungunya viral arthritic disease in humans

**DOI:** 10.1101/2023.11.20.567688

**Authors:** James Chang, Rimjhim Agarwal, Fernanda H Côrtes, Calvin Ha, John Villalpando, Izabella N. Castillo, Rosa Isela Gálvez, Alba Grifoni, Alessandro Sette, Claudia M Romero-Vivas, Mark T Heise, Lakshmanane Premkumar, Andrew K Falconar, Daniela Weiskopf

## Abstract

Chikungunya virus (CHIKV) is a mosquito-borne re-emerging viral infection that can cause chronic chikungunya viral arthritic disease (CHIKVD), which is characterized by incapacitating arthralgia and inflammation that can last for months to years following infection. Despite CHIKV outbreaks occurring worldwide and several vaccines currently in development, immune responses to CHIKV in humans remains largely understudied, and targets of T cell response are currently unknown. In this study, we tested peripheral blood mononuclear cells (PBMCs) collected from patients diagnosed with CHIKV infection during the 2014-2015 outbreak in Colombia against pools of overlapping peptides sequentially spanning each CHIKV protein. Using high-resolution flow cytometry, we detected robust CHIKV-specific CD4^+^, but not CD8^+^ T cell responses, in these patients. Patients still experiencing disease symptoms six years after infection displayed significantly stronger CHIKV-specific CD4^+^ T cell responses against nsP1, nsP2 and E2 proteins, compared to patients that resolved the infection. CHIKV-specific CD4+ T cells in symptomatic patients displayed a significantly lower Th1 CD4^+^ helper T cell responses and were enriched within the Th17 CD4^+^ helper subset, identifying tumor necrosis factor-alpha (TNFα) as the predominantly produced cytokine. In conclusion, this study comprehensively characterizes the T cell response against CHIKV in humans during the chronic phase and provides insights into the role of T cells and possible treatment of CHIKVD.

## INTRODUCTION

Chikungunya virus (CHIKV) is a mosquito-borne alphavirus in the family *Togaviridae*. The viral genome consists of two open-reading frames (ORFs) and encodes four non-structural proteins (nsP1- 4) responsible for the viral replication machinery and five structural proteins (Capsid, E3, E2, 6K, E1)^1^. The capsid proteins form the viral core that encapsulates the genomic RNA, while the E2 and E1 proteins play a pivotal role in facilitating viral assembly^2^.

First identified in Tanzania in 1952, CHIKV led to recurrent outbreaks in tropical and sub-tropical regions worldwide, including Africa and Asia, before causing a major outbreak in 2005-2006 in La Réunion Island, where almost a third of the population was infected^3–7^. In 2013, CHIKV emerged in the Americas, with initial cases in the Caribbean, and subsequently spread to Central and South America in 2014, causing a huge epidemic the following year^8^. The spread of CHIKV was facilitated by the expanding geographic range of *Aedes* mosquitoes due to rising temperatures and increasing travel, posing a significant public health concern^9,10^. Since the beginning of 2023, there have been over 440,000 cases and over 350 deaths reported from CHIKV globally^11^. Some vaccines, particularly PXVX0317 and VLA1553, have shown promising results in Phase II and III clinical trials, respectively; however, no vaccines have yet been licensed for global use^12,13^.

The acute phase of CHIKV infection is characterized by the sudden onset of febrile illness, maculopapular rash, and joint and muscle pain. Although the majority of infected individuals fully recover within 7-14 days, an estimated 30-60% of them experience incapacitating arthralgia that can persist for months to years^14–18^. Chronic infection predominantly affects small joints such as the phalanges and wrists and large joints such as ankles and shoulders^19^. Current therapeutic approaches primarily involve non-steroidal anti-inflammatory drugs (NSAIDs) and disease-modifying anti-rheumatic drugs (DMARDs); however, the efficacy of the treatment greatly varies between individuals^20^.

At present, the underlying mechanisms responsible for the development of chronic CHIKV symptoms remain elusive. In adult mice, CHIKV RNA could be detected 60-90 days post-infection in joint tissues, and in joints, muscle, and lymphoid tissue of rhesus macaques for weeks after infection^21–23^. A few studies have shown the presence of viral RNA and antigen using immunohistochemistry in synovial tissue of one patient 12 months post-infection and in muscle tissue biopsy specimen of another patient three months post-infection^24,25^. However, a more comprehensive analysis of synovial fluid via qRT-PCR and mass spectrometry in 38 patients showed no evidence of viral RNA or proteins 22 months post-infection^26^. However, multiple animal and human studies have revealed that joints are commonly infiltrated with macrophages and T cells, suggesting that chronic CHIKVD is immune-mediated^22,24,25,27–29^.

T cells play a pivotal role in combating viral infections. However, the targets of T cells and their possible role in CHIKV pathogenesis are not adequately investigated. Some studies have shown accumulation of activated CD8^+^ T cells in muscle and joint tissue biopsy samples collected from chronic patients^24,25^. Furthermore, a recent study suggests CHIKV can establish persistent infection in mice by evading CD8^+^ T cell responses^30^. Other studies have shown that CD4^+^ T cells are the primary mediators of joint inflammation and swelling in mice, which can be treated by suppressing T cell responses^31,32^. However, few studies have been performed to elucidate the function of T cells in chronic CHIKV infection in humans. As such, one study reported the presence of CHIKV-specific T cells one-two years after the La Réunion outbreak, but no difference was observed in patients still experiencing chronic disease compared to patients who resolved the disease^33^. However, since symptoms can persist for longer durations, it is essential to determine the involvement of T cells in the late chronic stages of the disease.

Here, using an *ex vivo* T cell stimulation assay, we show that CD4^+^ T cells play a pathogenic role in chronic CHIKVD in humans. PBMCs collected from patients infected during the 2014-2015 CHIKV epidemic in Colombia were stimulated with CHIKV-specific peptide megapools. The magnitude and functionality of CHIKV-specific CD4^+^ and CD8^+^ T cells were determined in an activation-induced Marker (AIM) and intracellular staining (ICS) assay, respectively. Interestingly, we found that chronic individuals have a significantly higher percentage of CHIKV-specific memory CD4^+^ T cells almost six years post-infection compared to recovered individuals, while CD8^+^ T cells were hardly detected in either group. Most responses in chronic patients were directed against nsP1, nsP2, and E2. Recovered individuals exhibited a significantly higher population of CCR6^-^ CXCR3^+^, which has been used to define Th1 T helper phenotype while CHIKV-specific CD4^+^ T cells in chronic individuals exhibited an enriched CCR6^+^ CXCR3^-^ population, which has been associated with Th17 T helper subset^34^. The majority of these cells produce a single cytokine, with TNFα being the most highly produced cytokine in the chronic group and IFNγ being the most highly produced in the recovered group. Overall, our work expands our current understanding of the pathogenesis of chronic CHIKV infection and provides an insight into possible therapeutic options.

## RESULTS

### Study donor cohorts

To characterize CHIKV-specific memory T cell targets, we enrolled 39 patients from Colombia who had been clinically diagnosed with CHIKVD, using an epidemiological diagnosis criterion. Infections were confirmed by the presence of CHIKV-IgG antibodies. Accordingly, the cohort has been divided into CHIKV seropositive (n=31) and seronegative (n=8) donors (Figure S1A). As shown in Table 1, both sexes were represented [28:3 (F:M) in the CHIKV seropositive cohort and 3:5 (F:M) in the CHIKV seronegative cohort]. At the time of sample collection, the average age of CHIKV-seropositive donors was 41 years, and the average age of seronegative donors was 38 years. This is consistent with demographics previously reported for gender and age groups affected^35^. On average, blood donations for both groups were obtained six years post-infection.

**Table 1:**
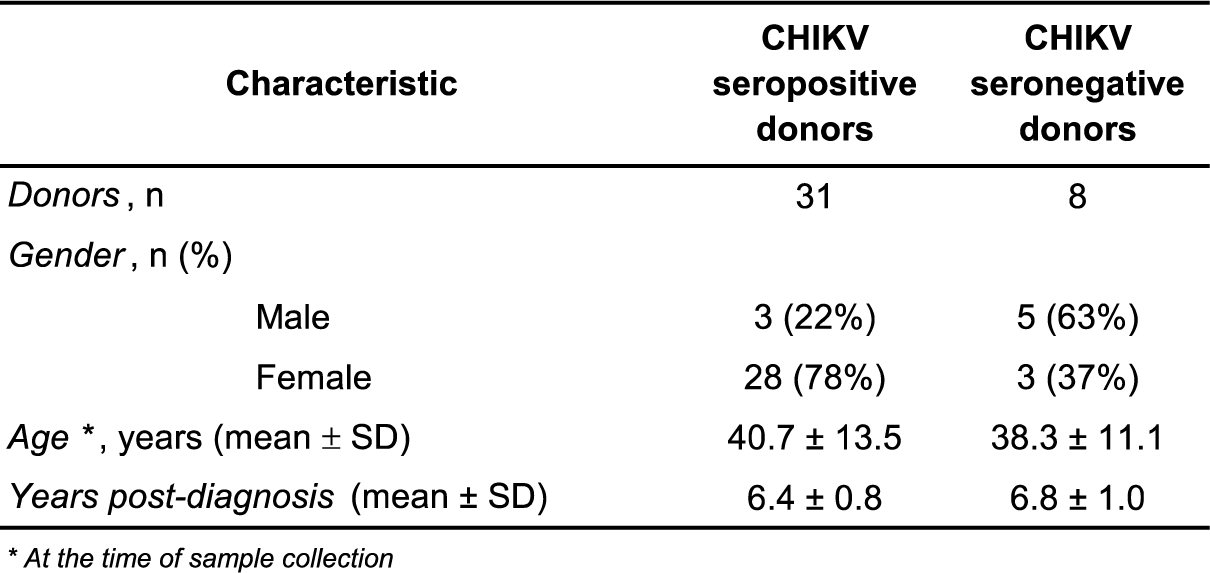
Characteristics of the donor cohort.

| Characteristic | CHIKV seropositive donors | CHIKV seronegative donors |
| --- | --- | --- |
| Donors, n | 31 | 8 |
| Gender, n (%) |  |  |
| Male | 3 (22%) | 5 (63%) |
| Female | 28 (78%) | 3 (37%) |
| Age *, years (mean ± SD) | 40.7 ± 13.5 | 38.3 ± 11.1 |
| Years post-diagnosis (mean ± SD) | 6.4 ± 0.8 | 6.8 ± 1.0 |

\* At the time of sample collection

### CHIKV-specific T cell responses

The frequency of CHIKV-specific T cell responses was determined using the previously described flow cytometry activation-induced marker (AIM) assay^36^. Specifically, CHIKV-specific CD4^+^ T cells were identified by upregulation of OX40^+^ and CD137^+^ while CHIKV-specific CD8^+^ T cells were identified by upregulation of CD69^+^ and CD137^+^. To determine immuno-dominance, we measured CHIKV-specific T cell responses against overlapping synthetic peptide sequences from each of the five structural proteins (CP, E3, E2, 6K, E1) and four non-structural proteins (nsP1, nsP2, nsP3, nsP4) (Figure 1A). Overall, CHIKV-specific CD4^+^ T cells were detected in 87% (27/31) of CHIKV-seropositive donors with responses directed against an average of three proteins (Figure 1B). In contrast, CHIKV-specific CD8^+^ T cells were only detected in 13% (4/31) of donors with responses against an average of two proteins (Figure 1C). CHIKV-specific CD4^+^ and CD8^+^ T cell responses were not detected in the CHIKV-seronegative control donors (0/8) (Figure S1B). CD4^+^ T cell responses to structural proteins and non-structural proteins accounted for 55% (41/75) and 45% (34/75) of the total CHIKV-specific CD4^+^ T cell responses, respectively (Figure 1B). Reactivity against E1, E2, and nsP1 accounted for the majority of CHIKV-specific CD4^+^ T cell responses (56%, 42/75) (Figure 1D). In contrast, responses against nsP4, E3, and 6K were infrequent (5%, 4/75) (Figure 1D). Antigen size did not seem to affect the CD4^+^ T cell targeting preferences as some of the largest proteins, such as nsP3 and nsP4 were infrequently recognized while smaller proteins such as nsP1, E2 and E1 proteins were immuno-dominantly recognized (Figure 1E).

**Figure 1:**
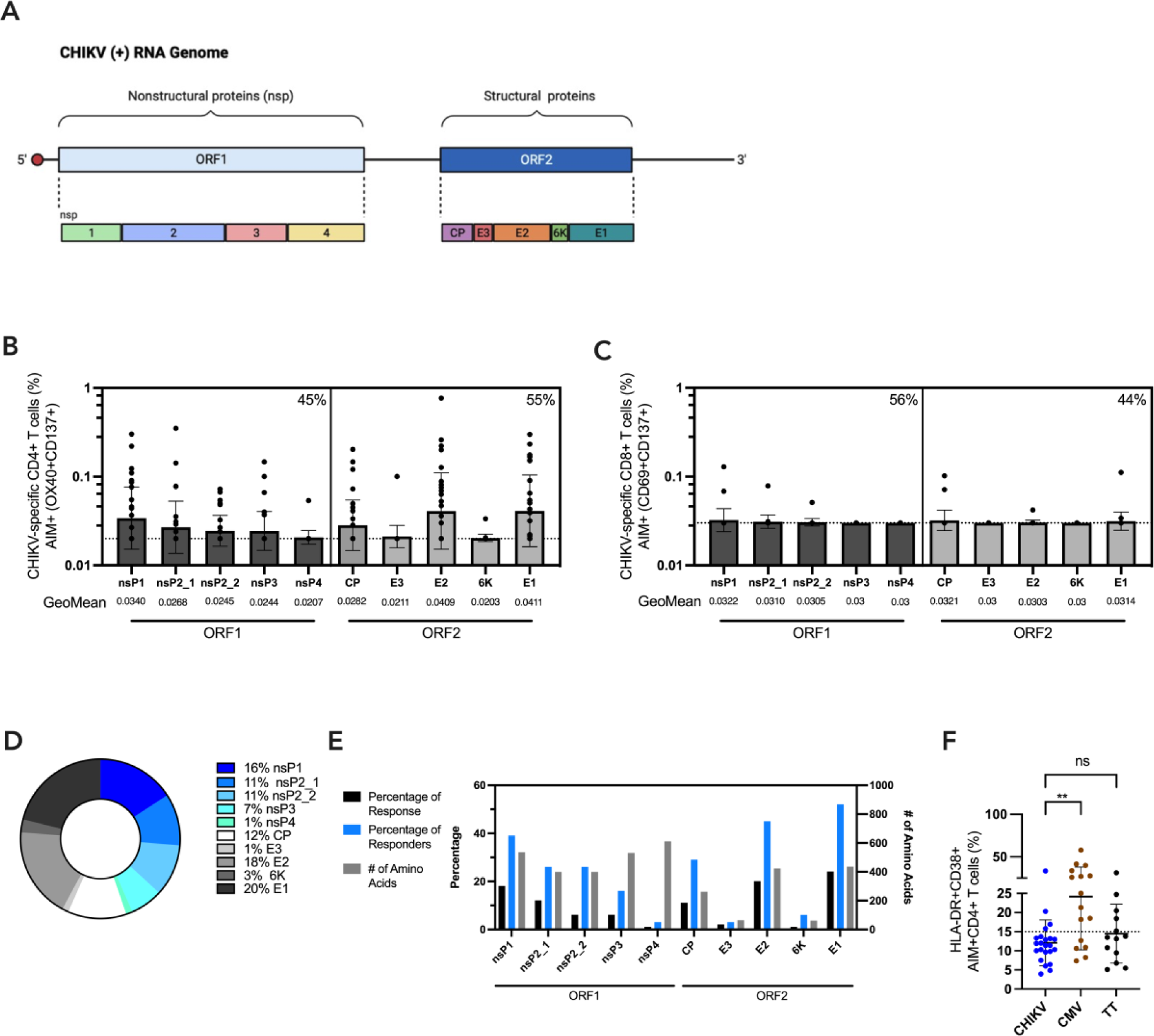
CHIKV-specific T cell responses in CHIKV-seropositive donors **(A)** Schematic representation of CHIKV genome comprised of non-structural proteins (nsP1-4) and structural proteins (CP, E3, E2, 6K, E1). **(B)** CHIKV-specific CD4^+^ T cells quantified by AIM (OX40^+^CD137^+^) after 24-hour stimulation with CHIKV pool per protein peptide megapools (MPs) in CHIKV-seropositive donors. **(C)** CHIKV-specific CD8^+^ T cells quantified by AIM (CD69^+^CD137^+^) after 24-hour stimulation with CHIKV pool per protein MPs in CHIKV-seropositive donors. **(D)** Frequency of AIM+ CD4^+^ T cell responses against all CHIKV proteins. **(E)** Summary of AIM+ CD4^+^ T cell responses per CHIKV protein and protein length. Black bars represent the percentage of responses *(n* = 75); blue bars represent the percentage of responders (*n* = 31); gray bar represents the number of amino acids present in each CHIKV protein. **(F)** Graph displaying CHIKV (blue dots), CMV (brown dots), and TT-specific (black dots) CD4^+^ T cell responses associated with recent activation. Recent activation was evaluated as the percentage of HLA-DR^+^ CD38^+^ from AIM+ (OX40^+^CD137^+^) CD4^+^ T cells. Data were analyzed for statistical significance using a Mann-Whitney *U* test. (p>0.05 = non-significant (ns), p<0.05 *, p<0.01 **).

Finally, we assessed if CHIKV-specific T cell responses reflected recent exposures to antigens. In previous studies, the expression of HLA-DR and CD38 activation markers has been linked to recent *in vivo* activation^37,38^. Markers for recent CD4^+^ T cell activation were measured against CHIKV, Cytomegalovirus (CMV), and tetanus toxoid (TT) derived peptide pools. (Figure 1F). CMV (a latent virus) represents frequent antigen exposure through chronic immune activation while TT represents infrequent antigen exposure, usually occurring through vaccine boosters every 10 years^39^. CHIKV- specific CD4^+^ T cell frequencies greater than 0.1% were evaluated for recent activation markers HLA-DR and CD38. The magnitude of recently activated CHIKV-specific CD4^+^ T cells was significantly less than CMV response (p = 0.0062), and more comparable to TT, thus closer resembling an infrequent rather than chronic exposure.

Overall, we identified a dominance of CHIKV-specific CD4^+^ T cell responses, predominantly against epitopes on the E1, E2, and nsP1 proteins, while CHIKV-specific CD8^+^ T cell responses were negligible in the chronic phase.

### CHIKV-specific CD4^+^ T cell responses in symptomatic and recovered individuals

It has previously been reported that CHIKV infection can lead to chronic infection, lasting months to years after infection^15^. Accordingly, we aimed to analyze if there was a difference in T cell responses in individuals currently still symptomatic versus recovered six years post-CHIKV infection. The clinical profile at the time of diagnosis for both groups is shown in Table 2. Patients that still display symptoms of arthralgia were categorized as “symptomatic” while patients without symptoms were categorized as “recovered”. Symptomatic patients had a slightly higher mean age (49.5 ± 11.6 years) compared to recovered patients (31.4 ± 6.4 years) at the time of sample collection (Table 2).

**Table 2:** Comparison of demographics and clinical symptoms between symptomatic and recovered individuals who are CHIKV-seropositive.

| Characteristic | Symptomatic | Recovered |
| --- | --- | --- |
| *Donors , n* | 17 | 14 |
| *Gender , n (%)* |  |  |
| Male | 2 (12%) | 2 (14%) |
| Female | 15 (88%) | 12 (86%) |
| *Age \*, years (mean ± SD)* | 49.5 ± 11.6 | 31.4 ± 6.4 |
| *Years post-infection (mean ± SD)* | 5.8 ± 0.5 | 6.9 ± 0.6 |
| *Current symptoms* |  |  |
| Athralgia** (%) | 100 | 0 |
| *Symptoms at the time of infection* |  |  |
| Fever (%) | 100 | 100 |
| Headache (%) | 100 | 100 |
| Non-purulent conjunctivitis (%) | 6 | 0 |
| Retrorbital pain (%) | 12 | 7 |
| Diarrhea (%) | 6 | 21 |
| Asthenia (%) | 65 | 14 |
| Myalgia (%) | 100 | 79 |
| Arthralgia (%) | 100 | 86 |
| Exanthema (%) | 75 | 57 |
| Edema lower limb and/or other parts of the body (%) | 76 | 21 |

\* At the time of sample collection
\*\* Includes pain in joints and upper and lower limbs, swelling in limbs, and muscle pain

We combined the response to all tested CHIKV megapools in symptomatic and recovered donors and found a significantly higher frequency of CHIKV-specific CD4+ T cells in symptomatic donors as opposed to those who had recovered (p < 0.0001) (Figure 2A). Next, we wanted to identify the specific CHIKV proteins that differed significantly between the two groups. The frequency of CHIKV-specific CD4^+^ T cell responses to nsP1, nsP2_1 and E2 (nsP1: p = 0.0475, nsP2_1: p = 0.0152, E2: p = 0.0269) were significantly higher in symptomatic individuals compared to recovered (Figure 2B). The frequency of CD4^+^ T cell responses against CP and E1 were higher in the symptomatic group but were not statistically significant (CP: p = 0.111, E1: p = 0.217). Additionally, on average, CHIKV-specific CD4^+^ T cells from symptomatic individuals responded to a greater number of CHIKV proteins than from recovered individuals (3.9 proteins in symptomatic and 1.8 proteins in recovered). Overall, immunodominance of CHIKV protein-specific responses detected were comparable between symptomatic and recovered individuals, with most responses against E1 and E2, followed by nsP1 and lastly nsP2 and CP (Figure 2C). We continued further analysis in donors that were positive in the AIM assay, based on the expression of OX40^+^CD137^+^. The magnitude of recently activated CHIKV-specific CD4^+^ T cells, based on the expression of HLA-DR and CD38, did not significantly differ between the symptomatic and recovered groups (Figure 2D).

**Figure 2:**
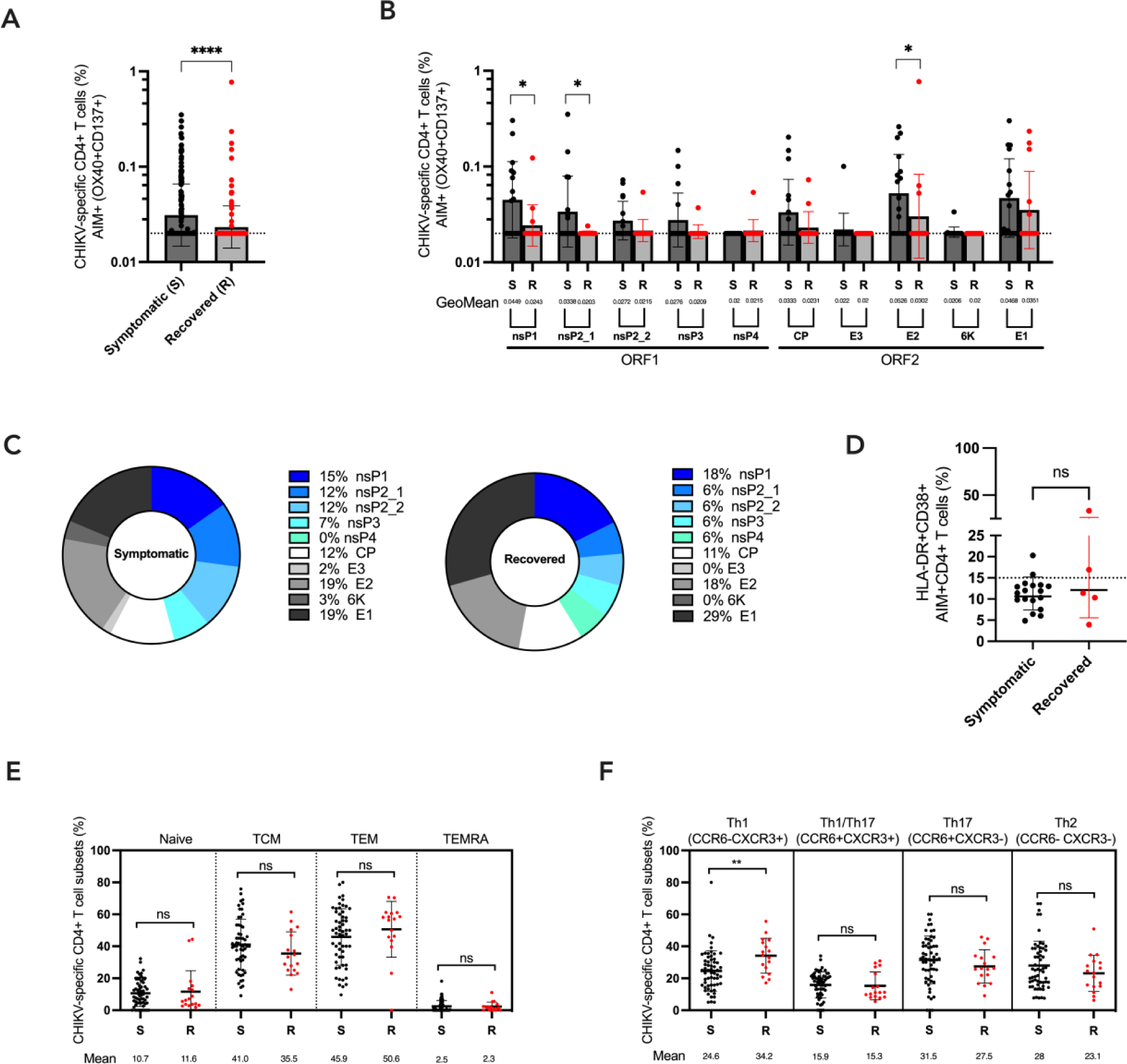
Immunodominance, recent activation, and memory phenotype of CHIKV-specific CD4^+^ T cells in symptomatic and recovered donors. (A) A combined CHIKV-specific CD4^+^ T cell response is shown to all CHIKV megapools in symptomatic (black dots) and recovered (red dots) donors. **(B)** CHIKV-specific CD4^+^ T cell response is shown per protein in symptomatic (black dots) and recovered (red dots) donors against each of the four non-structural (nsP1, nsP2_1, nsP2_2, nsP3, nsP4) and five structural proteins (CP, E3, E2, 6K, E1) **(C)** Pie charts display the distribution of CHIKV-specific CD4^+^ T cell responses per protein for symptomatic (*n* = 17) and recovered (*n* = 14) donors. **(D)** Comparison of the percentage of CHIKV-specific CD4^+^ T cells with expression of recent activation markers (HLA-DR^+^CD38^+^) in symptomatic (*n* = 17) and recovered (*n* = 5) donors, where AIM+ responses > 0.1% were used to evaluate recent activation. **(E)** CHIKV-specific CD4^+^ T cell memory subsets in the symptomatic (black dots) and recovered (red dots) donors have been defined based on the expression of CCR7 and CD45RA in AIM+ CD4^+^ T cells as: T naive (CCR7^+^ CD45RA^+^), TCM (T central memory; CCR7^+^ CD45RA^-^), TEM (T effector memory; CCR7^-^ CD45RA^-^) and TEMRA (T effector memory re-expressing CD45RA; CCR7^-^ CD45RA^+^). **(F)** CHIKV-specific CD4^+^ T helper cell (Th) subsets have been defined based on the expression of CXCR3 and CCR6 in AIM+ CD4^+^ T cells in symptomatic (black dots) and recovered (red dots) donors. Cells have been defined as Th1, Th1/Th17, Th17 or Th2 subsets. CHIKV-specific CD4^+^ T cells were analyzed for statistical significance using Mann-Whitney *U* test (A, B, D, E and F, p>0.05 = non-significant (ns), p<0.05 *, p<0.01 **, p<0.001***, p<0.0001****).

Next, we evaluated the memory phenotype of CHIKV-specific CD4^+^ T cells. The majority of CHIKV- specific CD4^+^ T cell memory subsets in both groups were CCR7^+^CD45RA^-^ central memory (TCM) and CCR7^-^CD45RA^-^ effector memory (TEM), and did not significantly differ between both groups (Figure 2E).

Finally, we evaluated CHIKV-specific CD4^+^ T helper cell (Th) subsets for symptomatic and recovered patients (Figure 2F), using CCR6 and CXCR3 markers, which have been used previously to define Th subsets^34^. Interestingly, recovered donors had a significantly higher proportion of CCR6^-^CXCR3^+^ CD4^+^ T cells (Th1) compared to symptomatic patients (p = 0.0046) (symptomatic mean: 24.6%, recovered mean: 34.2%), while the proportion of CCR6^+^CXCR3^-^ CD4^+^ T (Th17) cells was enriched in symptomatic patients (symptomatic mean: 31.5%, recovered mean: 27.5%). Interestingly, the proportion of CCR6^+^CXCR3^+^ CD4^+^ T cell population (Th1/Th17) (symptomatic mean: 15.9%, recovered mean: 15.3%) and the proportion of CCR6^-^CXCR3^-^ CD4^+^ T cells population (Th2) (symptomatic mean: 28.0%, recovered mean: 23.1%) was comparable in both groups (Figure 2F).

While the breadth and the magnitude of response were different in between symptomatic and recovered responders, we did not observe any significant differences in immunodominance, recent activation or memory phenotype of CHIKV-specific CD4^+^ T cells. Interestingly, we did observe a significant higher frequency of the Th1 subset in the recovered compared to the symptomatic group.

### Cytokine profile of CHIKV-specific CD4^+^ T cell responses

To determine the functional profiles of CHIKV-specific CD4^+^ T cells, we performed intracellular staining for cytokines (IFN*γ*, IL-2, TNFα, IL-17A, IL-4, IL-10), granzyme B (GzB) and CD40L (iCD40L) responses in donors that presented a positive response in the AIM assay (OX40^+^CD137^+^). IFN*γ*, IL- 2, TNFα, GzB, and IL-10 expression were reliably detected against nsP1, nsP2_1, CP, E2 and E1, while IL-4 and IL-17A expression was not detected (Figure S3). The cytokine responder profile for each CHIKV protein is shown in supplemental figure 3. TNFα was the most frequently expressed cytokine across all CHIKV proteins screened (produced in 94% of donors screened). nsP1, E1 and E2 induced IFNγ, IL-2, and TNFα expression in the greatest number of responders (Figure S3).

For proteins that elicited significant CHIKV specific CD4^+^ T cell responses in the AIM assay (nsP1, nsP2_1, and E2) between the symptomatic and the recovered donors, we then evaluated the multifunctional profile of CHIKV-specific CD4^+^ T cells in symptomatic donors. 71.7% of CHIKV-specific CD4^+^ T cells in symptomatic donors produced a single cytokine, 22.6% produced double cytokine, and 6.9% produced triple cytokine. None of the cells produced four cytokines (Figure 3A). The multifunctionality of CD4^+^ T cell response was also assessed against CMV as a model for chronic infection (Figure 3B). CMV induced significantly higher expression of IFNγ > IL-2 > TNFα single producers. Single-cytokine-producing AIM+ CD4^+^ T cells were compared between CHIKV and CMV (Figure 3C), where we observed a significantly higher percentage of IFN*γ* and IL-2 producing CD4^+^ T cells in response to CMV, as compared to the combined response to nsP1, nsP2_1 and E2 CHIKV proteins (IFN*γ*: p = 0.0002, IL- 2: p = 0.0160). However, most single-cytokine producing cells in CHIKV were primarily TNFα producers (p = 0.0215), in comparison to in CMV. We also evaluated the multifunctional CHIKV-specific CD4^+^ T cell responses in recovered patients, which also primarily produce IFN*γ* (Figure S4). In summary, CHIKV-specific CD4^+^ T cell response in symptomatic patients was characterized by the dominant production of TNFα.

**Figure 3:**
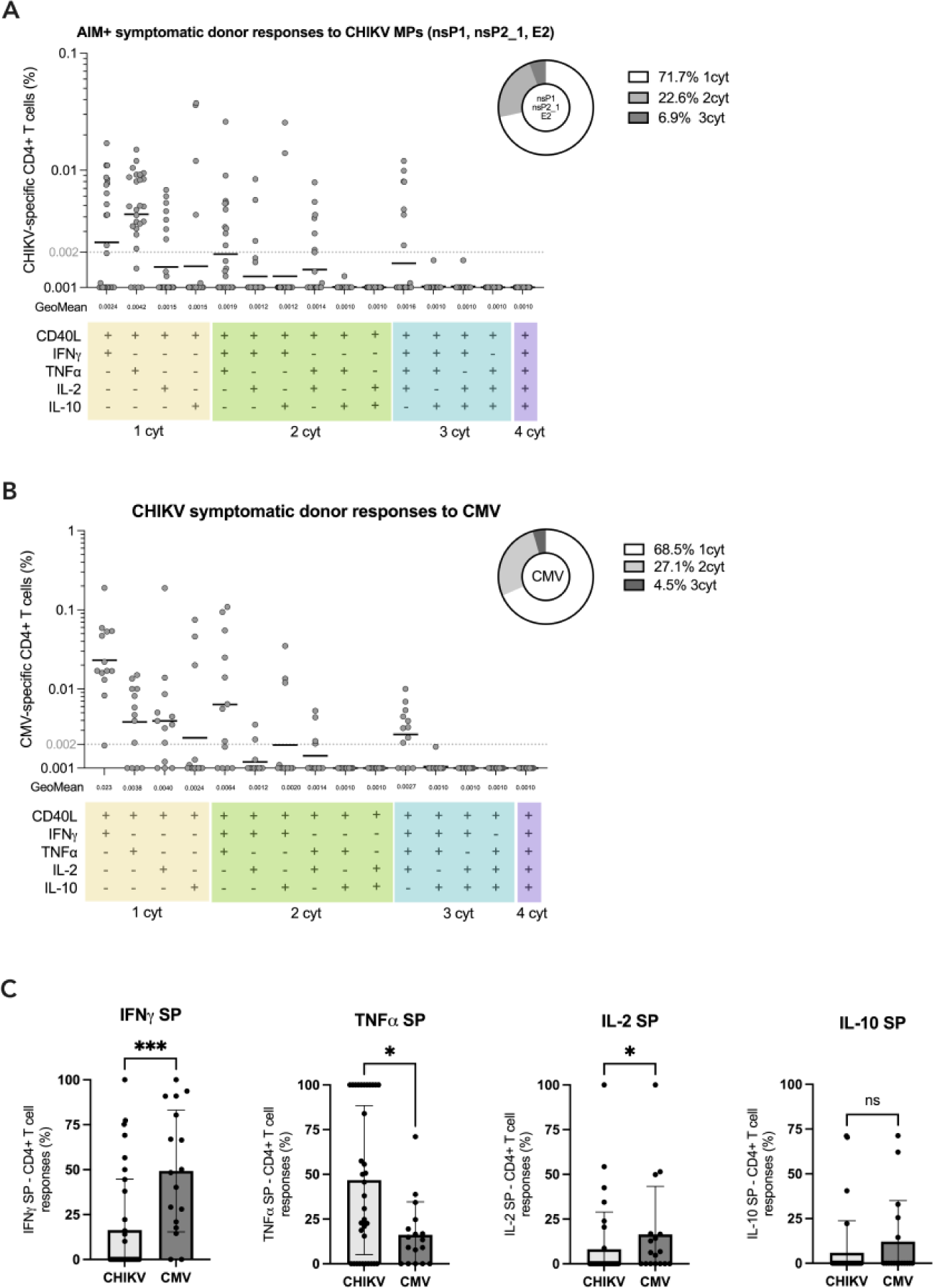
Memory CD4^+^ T cell multifunctional cytokine profile in AIM+ symptomatic donors **(A)** Quantification of cytokine+ CHIKV-specific CD4^+^ T cell (nsP1, nsP2_1, E2) multifunctional profile in symptomatic donors (*n* = 17), where the dotted black line indicates the limit of quantification (LOQ). Boolean gating was used to determine the multifunctional capacity of CHIKV-specific CD4^+^ T cells. The pie chart on the right shows the distribution of single, double and triple positive cytokine producing CHIKV-specific CD4^+^ T cells. **(B)** Quantification of the cytokine+ CMV-specific CD4^+^ T cell multifunctional profile in symptomatic donors (*n* = 17). **(C)** Comparison between single-cytokine producing CD4^+^ T cells in response to CHIKV (nsP1, nsP2_1, E2) and CMV for IFN*γ*, TNFα, IL-2 and IL-10 (SP = single producers). Data were analyzed for statistical significance using Mann-Whitney *U* test. (C, p>0.05 = non-significant (ns), p<0.05 *, p<0.01 **, p<0.001***).

## DISCUSSION

Our understanding of the role of T cells in chronic CHIKV infection is largely undefined and is primarily based on animal models. Previous studies have evaluated and detected CHIKV-specific memory T cells up to 24 months post-infection in humans^33^. In this study, using *ex vivo* stimulation of PBMCs collected from CHIKV-infected donors, we screened for CHIKV-specific T cell response against the five structural (CP, E3, E2, 6K, E1) and four non-structural (nsP1, nsP2, nsP3, nsP4) proteins. We found that over 87% of donors had detectable levels of CHIKV-specific memory CD4^+^ T cells in the periphery approximately six years post-infection, while CHIKV-specific CD8^+^ T cells were only detected in 13% of donors. This is consistent with previous work reporting diminished magnitude and breadth of CD8^+^ T cell responses in aged rhesus macaques with persistent infection^23^. In another study, the viraemia in spleen and joint tissue was reported to be indistinguishable in WT and CD8α^-/-^ mice, during the chronic phase of infection^30^. Therefore, our results further support the idea that CD8^+^ T cells do not play a significant role in controlling chronic CHIKV infection. Further studies are however required to elucidate whether CHIKV can evade CD8^+^ T cell responses and whether reactivating CHIKV-specific CD8^+^ T cells can aid in clearing or controlling infection during the chronic phase in humans.

In addition to a limited knowledge of the role of T cells, the target epitopes of T cells in CHIKV are not well understood, despite them being essential for vaccine design. By using megapools of synthetic peptides for each CHIKV protein, we observed no differences in the immunodominance of proteins that were recognized between symptomatic and recovered donors; however, interestingly, the magnitude of response to nsP1, nsP2 and E2 was significantly higher in symptomatic donors. All of the significantly higher recognized proteins play a critical role in viral replication. nsP1 encodes enzymes to cap viral RNA and anchors the replicase complex to the cell membrane, while E2 binds to receptors on the cell membrane^41^. E2 has further been shown to be a major target for neutralizing antibodies^42–45^. E2 and nsP1 have previously been shown to be recognized by circulating CHIKV-specific T cells 24 months post-infection, however, in opposition to our study, the authors did not observe any differences in responses between chronic and recovered donors^33^, although the authors have not studied the phenotype past 24 months of infection.

It has been previously demonstrated that CD4^+^ T cells contribute to joint swelling and inflammation in mice^31,32^. Here, we found a significantly higher magnitude of CD4^+^ T cell response against the nsP1, nsP2, and E1 CHIKV proteins in symptomatic donors compared to the recovered donors, which clearly implicates CD4^+^ T cells in the development of chronic disease. Thus, our results provide the first association of CHIKV-specific T cells in chronic CHIKV pathogenesis in humans

Previous studies have reported the presence of viral antigen or RNA in joint tissues of mice and non-human primates 60-90 days and up to three months after infection, respectively^21–23^. However, to our knowledge, viral antigen has only been detected as far as 12 months post-infection in humans^21^ and was no longer detected 22 months post-infection^26^. Our data describing the lack of recent activation markers in CHIKV-specific T cells also supports the absence of viral antigen in patients with chronic symptoms.

Another prevalent hypothesis in the field to explain CD4^+^ T cell pathogenicity is the idea of T cell-mediated autoimmunity, which is also a mechanism associated with rheumatoid arthritis (RA)^46–49^. Our findings are more in accordance with the latter hypothesis as we observed an enrichment of CCR6^+^CXCR3^-^, which are associated with Th17 cells, in the symptomatic group, while the recovered group exhibited a significantly higher CCR6^-^CXCR3^+^, which are associated with Th1 phenotype^34^.

Th17 cells have been implicated in RA, and even though we did not detect any production of IL- 17A, previous work have highlighted that IL-17A is not sufficient to exclude the role of Th17 cell in pathogenicity ^50–56^. Additionally, our results are consistent with previous work done in mice showing that absence of TNFα receptor improved inflammatory symptoms post-CHIKV infection^57^. TNFα has also been shown to be upregulated in RA patients and TNFα inhibitors have become a common treatment strategy for RA^58–60^, similar to what we observed in symptomatic individuals where 94% of symptomatic individuals produced TNFα, in response to either nsP1, nsP2 and E2. Interestingly, over 70% of CD4^+^ T cells in symptomatic individuals were monofunctional, which is concerning, since polyfunctional CD4^+^ T cell response has been associated with reduced disease severity in other RNA viral infections such as in influenza infection^61^. Therefore, our work suggests that Th1 cell type could potentially be protective in chronic infection, while monofunctional, TNFα producers and Th17 cells contribute to the development of disease. Additional studies are required to define specific T helper subsets in CHIKV infection and to elucidate the precise role of Th17 cells and Th1 cells in chronic CHIKV particularly in affected tissues such as joints and synovium, where Th17 cells and non-classical Th1 are often found^62^.

Overall, our work indicates that CD4^+^ T cells, which primarily produce TNFα upon recognition of nsP1, nsP2, and E2, contribute to the chronicity of CHIKV infection and potentially lead to the development of arthralgia. Therefore, TNFα inhibitors could potentially be repurposed to aid individuals with persistent arthritis-like symptoms, although, extensive research is needed to understand the underlying mechanism by which TNFα is preferentially produced by CD4^+^ T cells in CHIKV and if anti-TNFα therapy could be beneficial.

Strengths and limitations of this study

CHIKV has been shown to cause tissue damage and inflammation. While it has been suggested in various animal models that T cells are involved in pathogenesis, this is the first human study that demonstrates a strong association between CHIKV-specific CD4^+^ T cells and chronic disease. Due to limited access to human samples, we were unable to assess T cell responses in affected tissues and synovium, which could provide more insights into phenotypes of tissue-resident memory T cells. Additionally, due to the same reasons, we could not test T cell responses during the acute phase of infection or longitudinally to understand how magnitude and breadth of T cell reactivity is modulated over the course of infection. We are aware that more work is required to determine the association between different CD4^+^ T cell types in CHIKV and chronic disease development.

## Acknowledgements

We thank all members of Weiskopf and Sette lab for their helpful discussions.

## Funding

This work was funded by National Institute of Health (NIH) Contract No. 75N93019C00065 to A.S. and D.W.

## Competing Interests

D.W is a consultant for Moderna. The remaining authors declare no conflicts of interest.

## MATERIALS AND METHODS

### Study cohort

We enrolled 39 donors who had been diagnosed with CHIKV during the 2014-2015 epidemic in the study at our site in Colombia in 2021. The criteria of a positive CHIKV diagnosis included: 1) whether the signs and symptoms were compatible with CHIKV, 2) whether the patient resided in or visited a region where circulation of CHIKV had been detected using a RT-PCR assay, 3) whether the patient lived in the same household/neighborhood as other residents who had been diagnosed with CHIKV, 4) whether the infections with dengue virus, which often co-circulates with CHIKV and has similar clinical presentation, has been ruled out through laboratory tests such as ELISA and rapid tests. Not all patient samples were subjected to confirmatory laboratory tests. The epidemiological link based on the criteria mentioned above was considered sufficient to diagnose a case as CHIKV positive by a local medical professional. For the purpose of our studies, we confirmed infection by measuring the levels of CHIKV-IgG antibodies using the serum of all individuals.

At the time of enrollment in the study, all individual donors provided informed consent that their samples could be used for any future studies, including this study. All blood and serum samples were anonymized and were given an anonymous code number. In addition to collecting blood and serum samples, information regarding the date of their CHIKV diagnosis, the date of symptom onset, the date of sample collection, their age, their gender, their encounters with other diseases, and the specific symptoms at the time of CHIKV diagnosis and sample collection was also collected for each participant.

In the study cohort, the average age of the donors was 39.5 years and both sexes were represented [31:8, F:M]. Information regarding their symptoms at the time of CHIKV infection and the current status of symptoms is provided in Table 1 and Table 2.

### Study approval

All samples were collected under an approved IRB protocol (LJI# VD154).

### PBMC isolation

Whole blood samples were collected from CHIKV donors at our site in Colombia. Large volumes of the donors’ blood were collected in sterile bags which contained 3.27 g citric acid, 26.3 g NaH_2_- citrate and 2.22 g NaHCO3 per liter (Fresenius Kabi, Fresenius HemoCare, Brasil Ltd, Brasil) and then diluted by 50% v/v with RPMI-1640 media (31800-105, Gibco, USA) containing 0.2% w/v NaHCO3 and 34ml volumes were gentle layered into multiple sterile 50 ml tubes (430290, Corning, USA) containing 12 ml of sterile endotoxin-tested Ficoll-Paque PLUS (1.077 g/ml density) (GE Healthcare Biosciences, Sweden). After centrifugation at 500xg, the buffy-coat peripheral blood mononuclear cells (PBMCs) were collected from immediately above the Ficoll-Paque-plasma interface, diluted with 50% v/v RPMI 1640 medium and again centrifuged at 500 x g to pellet the PBMCs. These PBMCs were then re-suspended in large volumes of RPMI 1640 medium containing 5% v/v fetal calf serum (FCS) (F-0926, Sigma, USA) and centrifuged against at 500 x g and after a further wash their PMBCs were re-suspended in ice-cold 10% dimethyl sulfoxide (D-2650, Sigma, USA) in FCS at approximately 20 x 10^6^ PBMCs/ml and 1 ml aliquots contained in 1.5 ml polypropylene cryo-vials (5000-1020: Nalgene System 100, Thermo Scientific, USA) were slowly (-1°C/min) frozen down to -80°C in a isopropanol alcohol-walled freezing unit (Nalgene Mr. Frosty: C-1562 Sigma-Aldrich, USA) before being transferred to 25-vial boxes in liquid nitrogen.

### Peptide pools

Since only a limited number of full-length sequences of CHIKV polyprotein were available from NIAID Virus Pathogen Database and Analysis Resources (ViPR) database, we made two separate consensus alignments for structural and non-structural polyprotein sequences and retrieved 257 and 350 full-length CHIKV structural and non-structural polyprotein sequences from ViPR using the following query: Chikungunya virus, Gene product name: structural OR non-structural polyprotein, Remove duplicate sequences. Unresolved sequences were removed.

The number of sequences available varied as a function of geographic locations. To ensure balanced representation, the number of isolates by geographical region was limited to a maximum of 10. In total, 158 structural and 61 non-structural sequences were selected. For each polyprotein, sequences were aligned using MUSCLE, and consensus sequences were BLASTed to identify a representative isolate (GenBank ID: AQX78118.1 and AQX78116.1), using tools in ViPR. To account for variants to the consensus sequences, we additionally synthesized 53 and 71 amino acid variants with a frequency above 10% from structural and non-structural consensus sequences, respectively.

Based on the consensus sequence and the variants, we synthesized 15-mer peptides overlapping by 10 residues. Together, peptides that spanned the length of the CHIKV proteome were synthesized, resuspended in dimethyl sulfoxide (DMSO), and divided into 11 pools. Number of peptides per megapool are listed in Table S1. Since the nsP2 megapool was too large, it was split into two pools (nsP2_1 and nsP2_2) of 117 and 81 peptides, respectively.

This megapool approach has been used previously to simultaneously screen a large number of epitopes. In this approach, large numbers of different epitopes are solubilized, pooled, and re-lyophilized to avoid cell toxicity problems associated with high concentrations of DMSO typically encountered when pooling after a single solubilization step. These MPs have been used in several indications, including SARS-CoV-2^63^, allergies^64^, tuberculosis^65^, tetanus, pertussis^66,67^, and DENV for both CD4^+^ and CD8^+^ T cell epitopes^68,69^.

### Activation-induced cell marker (AIM) T cell assay

PBMCs were thawed in 10 mL of RPMI 1640 (Corning) supplemented with 5% human AB serum (GeminiBio), penicillin [100 IU/mL], streptomycin [100μg/mL] (GeminiBio), and 2 mM L-glutamine (Gibco), and in the presence of benzonase (20μL/10mL). PBMCs were then plated at 1 x 10^6^ cells per well in 96-well U bottom plates and stimulated with CHIKV-specific megapools (MPs) [1μg/mL]. An equimolar amount of DMSO was used for negative control. Stimulation with phytohemagglutinin (PHA, Roche) [1μg/mL], a combined CD4 and CD8 CMV MP, and a tetanus-toxoid (TT) MP were used for positive controls [1μg/mL]. After stimulation for 24h, cells were stained for detection of activation-induced markers and measured by spectral flow cytometry on Cytek Aurora (Cytek Biosciences, USA). The antibody panel utilized for surface staining CD4^+^ and CD8^+^ T cells is shown in Table S2.

Antigen-specific CD4^+^ T cells were measured as a percentage of AIM+ (OX40^+^ CD137^+^) CD4^+^ T cells and antigen-specific CD8^+^ T cells were measured as the percentage of AIM+ (CD69^+^ CD137^+^) CD8^+^ T cells. The stimulation index (SI) was calculated by dividing the percentage of stimulated samples by those of the DMSO control. A SI > 2 and limit of detection (LOD) > 0.02% after background subtraction (DMSO), was considered a positive response for antigen-specific CD4^+^ T cells. A SI > 3 and LOD > 0.03% after background subtraction (DMSO) was considered a positive for antigen-specific CD8^+^ T cells. Expression of HLA-DR and CD38 markers was used to measure *in vivo* recent activation of CHIKV-specific CD4^+^ T cell, as has been shown previously in SARS-CoV-2 infection and other common cold viral infections^37,70–72^. A representative gating strategy is shown in supplemental Figure S2A.

### Intracellular cytokine staining (ICS) T cell assay

All samples that were defined to be positive in the AIM assay were subjected to further analysis in the ICS assay. PBMCs were plated at 1x10^6^ cells per well in 96-well U bottom plates and blocked with anti-CD40 Ab for 15 min at 37°C. Cells were then stimulated with CHIKV-specific MPs [1μg/mL] for 18h. An equimolar amount of DMSO was used for negative control, and stimulation with PHA [1μg/mL] and a combined CD4 and CD8 CMV MP [1ug/ml] were used for positive control. After 14h, Golgi-Plug and Golgi-Stop were added to the culture, in addition to anti-CD69 and anti-CD137 Abs. Cells were then washed, incubated with BD human FC block, and stained with LIVE/DEAD marker in the dark for 15 min. After incubation, cells were washed, surface stained in the dark for 30 min at 4°C, and then fixed with 1 % of paraformaldehyde (Sigma-Aldrich, St. Louis, MO). Subsequently, cells were permeabilized and stained with intracellular antibodies in the dark for 30 min at RT. All samples were acquired on Cytek Aurora. Antibodies used in this assay are listed in Table S3. A representative gating strategy is shown in supplemental Figure S2B.

Antigen-specific CD4^+^ T cells in the ICS assay were defined using iCD40L and cytokine-specific markers. These were then background subtracted (DMSO) and subjected to boolean analysis using FlowJo 10.8.1 to determine their multifunctional profile. For the ICS analysis, limit of sensitivity (LOS) > 0.002% was considered positive.

### Serologic assays

Site-specifically biotinylated CHIKV E1 DIII and Halo-tag control antigens were coupled to unique MagPlex®-Avidin microspheres at a concentration of 5μg of antigen per 10^6^ beads in assay buffer (1% BSA + phosphate-buffered saline (PBS), pH 7.4) for 1 hour at 37°C with shaking at 700 rpm as described before^73^. Antigen coupled beads were washed and aliquoted 2,500 beads per antigen per well into a 96-well assay plate. Heat-inactivated (56°C for 30 min) human serum samples diluted at 1:500 in assay buffer were incubated with beads for 1 hour at 37°C with shaking at 700 rpm. After washing the beads with assay buffer, PE conjugated goat anti-human IgG Fc secondary Ab was added at 6μg/ml (Southern Biotech, catalog: 2014-09) and incubated for 1 hour at 37°C with shaking at 700 rpm. Beads were washed and resuspended in 100μL assay buffer for fluorescence analysis using the Luminex 200 system.

### Quantification and statistical analysis

Data and statistical analyses were done in FlowJo 10.8.1 and GraphPad Prism 9 (La Jolla, CA) unless otherwise stated. The statistical details of the experiments are provided in the respective figure legends. Data plotted in linear scale were expressed as Mean ± Standard Deviation (SD). Data plotted in logarithmic scales were expressed as Geometric Mean ± Geometric Standard Deviation (SD). Mann-Whitney or Wilcoxon tests were applied for unpaired or paired comparisons, respectively. Kruskal-Wallis test adjusted with Dunn’s test for multiple comparisons was used to compare multiple groups.

## Notes

### Competing Interest Statement

D.W. is a consultant for Moderna. The remaining authors declare no competing interests.

