## Supplemental Material for "Chikungunya-virus-specific CD4^+^ T cells are associated with chronic chikungunya viral arthritic disease in humans"

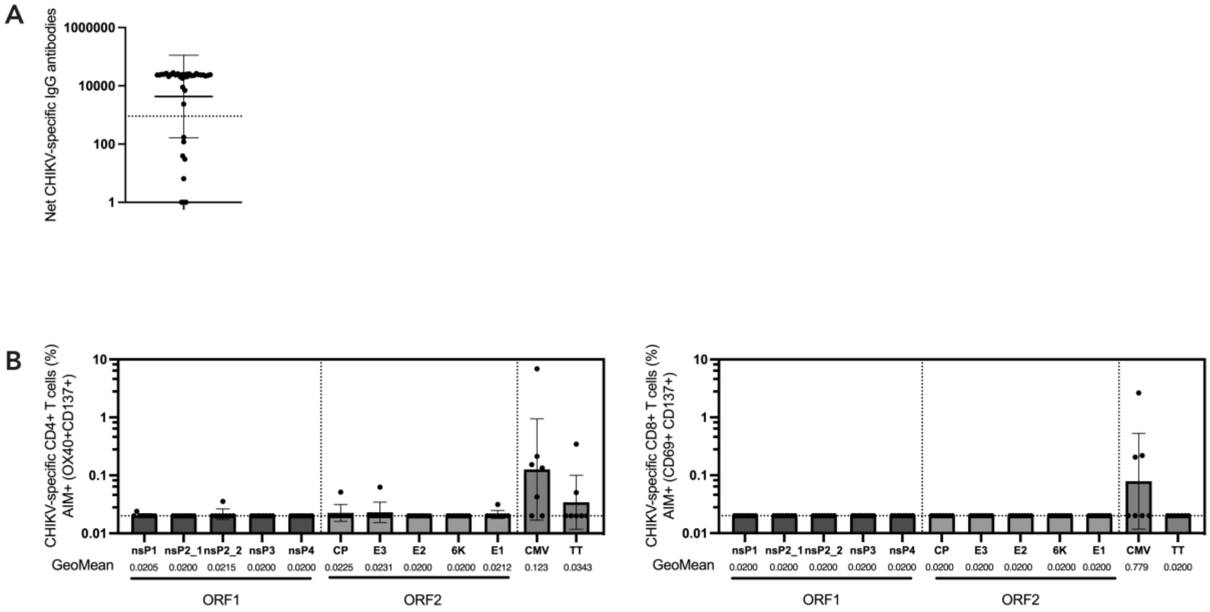


**Fig S1. Assessment of CHIKV-specific IgG antibodies and T cells**
(**A**) CHIKV-specific IgG measured in CHIKV cohort (n = 39). A threshold of 907 (dotted line) was used to determine seropositivity and to confirm CHIKV infection. **(B)** CHIKV-specific $CD4^{+}$ and $CD8^{+}$ T cells against CHIKV structural (CP, E3, E2, 6K, E1) and non-structural (nsP1, nsP2_1, nsP2_2, nsP3, nsP4) MPs, CMV and TT (for controls) in CHIKV seronegative donors (*n* = 8).

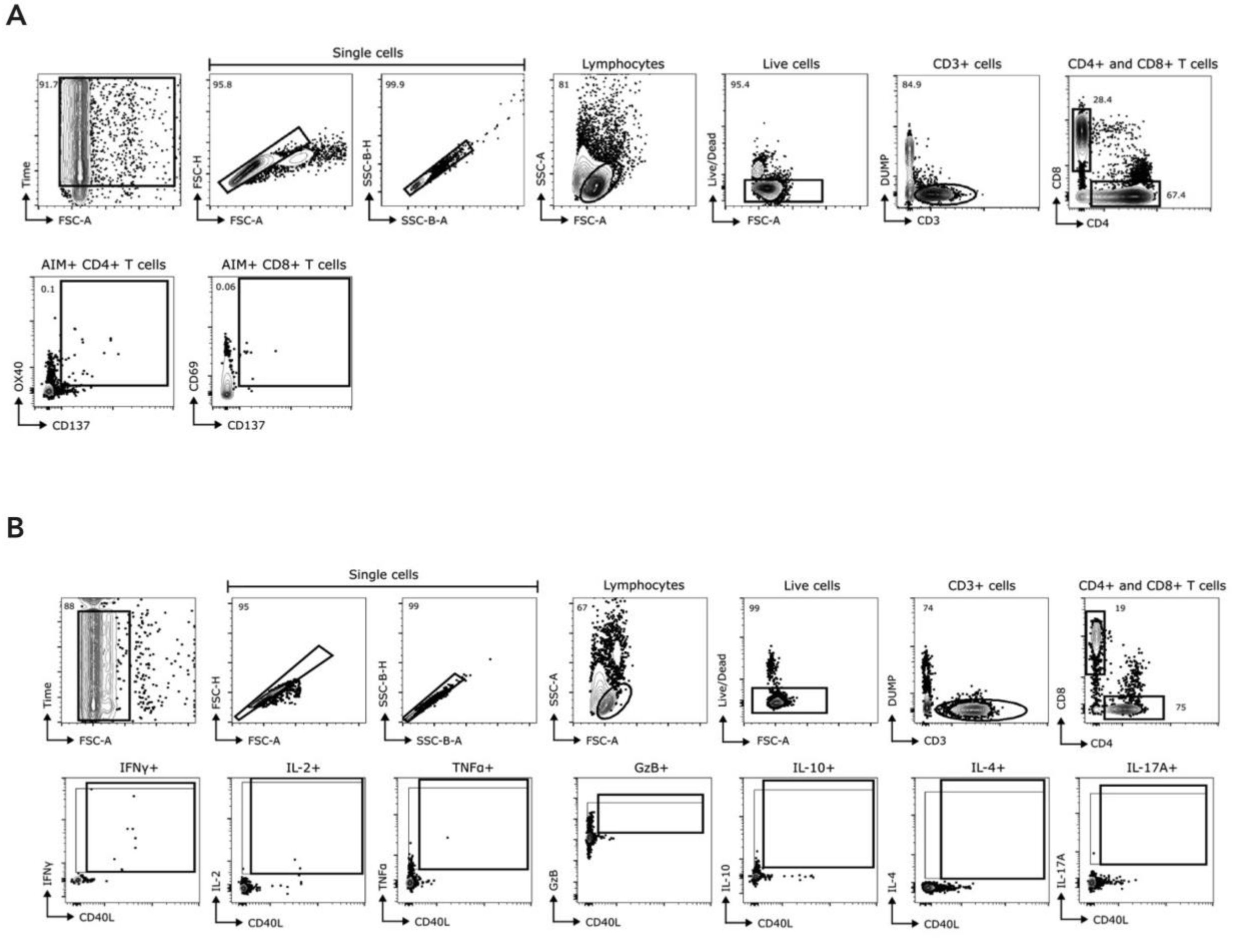


**Fig S2. Representative gating strategies for AIM and ICS assays**
(A) Representative gating strategy to define CD3$^+$CD4$^+$ and CD3$^+$CD8$^+$ cells by AIM assay. **(B)** Representative gating strategy to define CD3$^+$CD4$^+$ and CD3$^+$CD8$^+$ cells and intracellular CD40L$^+$cytokine$^+$ cells in ICS staining of cytokine (IFNγ, IL-2, TNFα, IL-17A, IL-4), granzyme B (GzB), and intracellular CD40L against CHIKV non-structural (nsP1-4) and structural (CP, E3, E2, 6K, E1) proteins. AIM+ CD4$^+$ responses LOD > 0.02% ($n$ = 75) were evaluated for cytokine production.

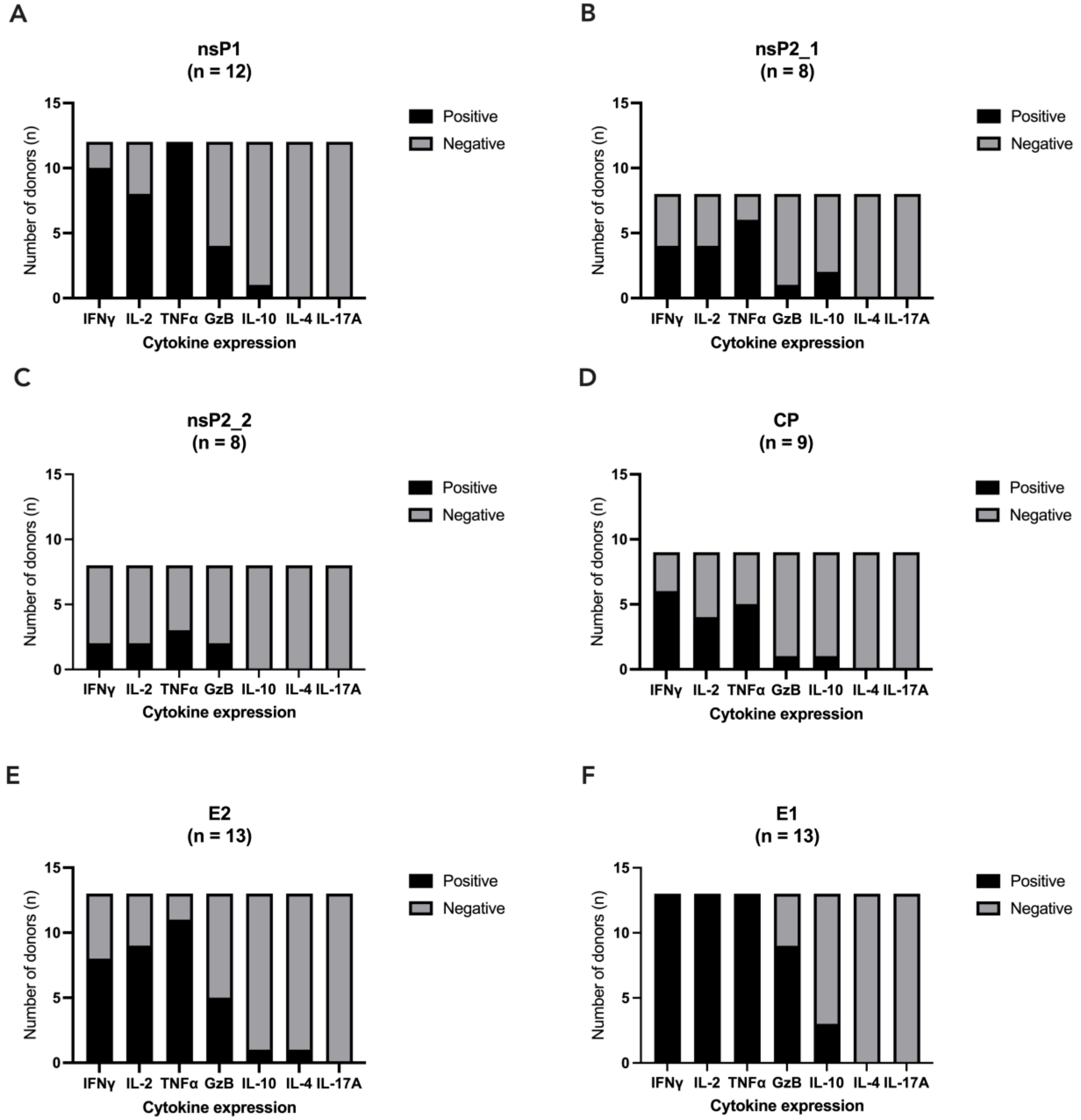


**Fig S3: Cytokine responder profile per CHIKV protein**

**(A-F)** The proportion of cytokine expression among AIM+ donors against individual CHIKV protein. n represents the number of donors tested. The black bars represent the number of donors that tested positive and the gray bars represent the number of donors that tested negative for specific cytokine. CHIKV proteins that were screened more than five times are shown.

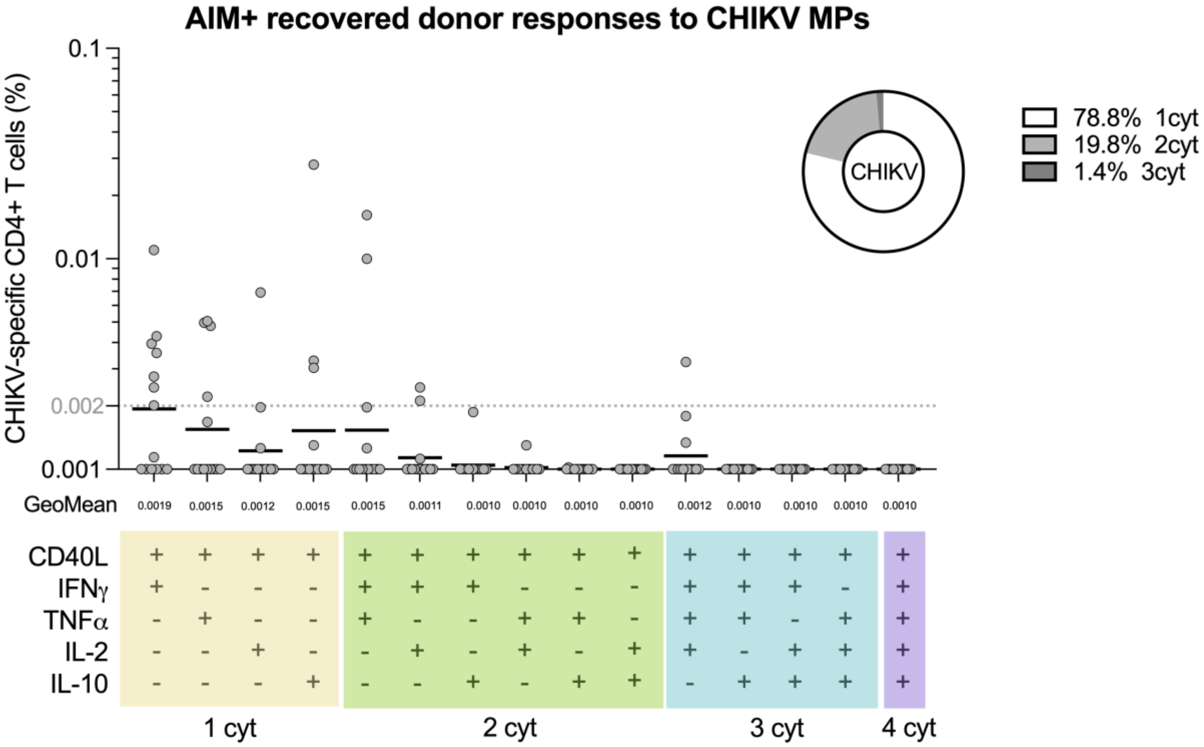


**Fig S4: Memory CD4⁺ T cell multifunctional cytokine profile in AIM+ recovered donors.** Quantification of cytokine+ CHIKV-specific CD4⁺ T cell (nsP1-4, CP, E2, E1) in recovered donors ($n$ = 8).

### SUPPLEMENTAL TABLES

**Table S1. Number of overlapping 15-mer peptides per CHIKV protein**

| Protein | # of peptides per protein |
| --- | --- |
| CP | 68 |
| E3 | 25 |
| E2 | 124 |
| 6K | 23 |
| E1 | 113 |
| nsP1 | 133 |
| nsP2_1 | 117 |
| nsP2_2 | 81 |
| nsP3 | 153 |
| nsP4 | 155 |
| Total | 992 |

### Table S2: Reagents used for AIM assays

| Reagent | Clone (Source) | Catalog. No. | Dilution |
|---|---|---|---|
| Live/Dead Blue | (ThermoFisher) | L23105 | 1:1000 |
| CCR6-BUV496 | 11A9 (BD Biosciences) | 612948 | 1:200 |
| CXCR5-BV421 | J252D4 (BioLegend) | 356920 | 1:200 |
| CXCR3-BV605 | G025H7 (BioLegend) | 353728 | 1:200 |
| CCR7-BV711 | G043H7 (BioLegend) | 353228 | 1:200 |
| CD3-BUV395 | UCHT1 (BD Biosciences) | 563546 | 1:1000 |
| CD137-BUV737 | 4B4-1 (BD Biosciences) | 741861 | 1:50 |
| CD8-BUV805 | SK1 (BD Biosciences) | 612889 | 1:1000 |
| CD16-BV510 | 3G8 (BioLegend) | 302048 | 1:1000 |
| CD14-BV510 | 63D3 (BioLegend) | 367124 | 1:1000 |
| CD20-BV510 | 2H7 (BioLegend) | 302340 | 1:1000 |
| CD45RA-BV570 | HI100 (BioLegend) | 304132 | 1:200 |
| CD38-BV650 | HB-7 (BioLegend) | 356620 | 1:200 |
| PD-1-BV785 | EH12.2H7 (BioLegend) | 329930 | 1:200 |
| CD69-FITC | FN50 (BioLegend) | 310904 | 1:100 |
| CD4-cFluor b548 | SK3 (Cytek Biosciences) | R7-20043 | 1:200 |
| CD95-BB700 | DX2 (BD Biosciences) | 566542 | 1:200 |
| CD40L-PE-Dazzle594 | 24-31 (BioLegend) | 310840 | 1:200 |
| OX40-APC | Ber-Act35 (BioLegend) | 350008 | 1:100 |
| HLA-DR-APC-R700 | G46-6 (BD Biosciences) | 565127 | 1:200 |
| CD40 | HB14(Miltenyi Biotec) | 5210201190 | 1:133 |

### Table S3: Reagents used for ICS assays

| Reagent | Clone (Source) | Catalog. No. | Dilution |
|---|---|---|---|
| Live/Dead Blue | (Thermo Fisher) | L23105 | 1:500 |
| CD3-BUV395 | UCHT1 (BD Biosciences) | 563546 | 1:100 |
| CD8-BUV805 | SK1 (BD Biosciences) | 612889 | 1:200 |
| CD16-BV510 | 3G8 (BioLegend) | 302048 | 1:200 |
| CD14-BV510 | 63D3 (BioLegend) | 367124 | 1:200 |
| CD20-BV510 | 2H7 (BioLegend) | 302340 | 1:200 |
| CD45RA-BV570 | HI100 (BioLegend) | 304132 | 1:100 |
| CD4-cFluor b548 | SK3 (Cytek Biosciences) | R7-20043 | 1:25 |
| CD69-BV605 | FN50 (BD Biosciences) | 562989 | 1:100 |
| CCR7-PE-Cy7 | G043H7 (BioLegend) | 353226 | 1:100 |
| IL-4-BUV737 | MP4-25D2 (BD Biosciences) | 612835 | 1:200 |
| IL-17-BV785 | BL168 (BioLegend) | 512338 | 1:200 |
| IFNγ-FITC | 4S.B3 (Thermo Fisher) | 11-7319-82 | 1:500 |
| IL-2-BB700 | MQ1-17H12 (BD Biosciences) | 566405 | 1:200 |
| IL-10 -PE-Dazzle594 | JES3-19F1 (BioLegend) | 506812 | 1:200 |
| TNFα-eFluor450 | Mab11 (Thermo Fisher) | 48-7349-42 | 1:200 |
| Granzyme B-AF647 | GB11 (BD Biosciences) | 560212 | 1:100 |
| CD40L-PerCP-ef710 | 24-31 (Thermo Fisher) | 46-1548-42 | 1:50 |
| CD137-PE-Cy5 | 4B4-1 (BD Biosciences) | 551137 | 1:100 |
